# TLR2 signaling pathway combats *Streptococcus uberis* infection by inducing production of mitochondrial reactive oxygen species

**DOI:** 10.1101/809186

**Authors:** Bin Li, Zhixin Wan, Zhenglei Wang, Jiakun Zuo, Yuanyuan Xu, Xiangan Han, Vanhnaseng Phouthapane, Jinfeng Miao

**Author notes:** These authors contributed equally (Bin Li, Zhixin Wan).

## Abstract

Mastitis caused by *Streptococcus uberis* is a hazardous clinical disease in dairy animals. In this study, the role of Toll-like receptors (TLRs) and TLR-mediated signaling pathways in mastitis caused by *S. uberis* was investigated using mouse models and mammary epithelial cells (MECs). We used *S. uberis* to infect mammary glands of wild type, TLR2^−/−^ and TLR4^−/−^ mice and quantified the adaptor molecules in TLR signaling pathways, proinflammatory cytokines, tissue damage and bacterial count in mammary glands. When compared with TLR4 deficiency, TLR2 deficiency induced more severe pathological changes through myeloid differentiation primary response 88 (MyD88)-mediated signaling pathways during *S. uberis* infection. In MECs, TLR2 detected *S. uberis* infection and induced mitochondrial reactive oxygen species (mROS) to assist host control of secretion of inflammatory factors and elimination of intracellular *S. uberis*. Our results demonstrate that TLR2-mediated mROS have a significant effect on *S. uberis*-induced host defense responses in mammary glands as well as MECs.

**Author summary:** *S. uberis* contributes significantly to global mastitis and remains a major obstacle for inflammation elimination due to its ability to form persistent infection in mammary tissue. The Toll-like receptor (TLR) family plays a significant role in identifying infections of intracellular bacteria and further triggering inflammatory reactions in immune cells. However, the detailed molecular mechanism by which TLR is regulated, and whether MECs, as the main cells in mammary gland, are tightly involved in these processes is poorly understood. Here, we used *S. uberis* to infect mammary glands of wild type, TLR2^−/−^, TLR4^−/−^ mice and MECs to assess pathogenesis, proinflammatory cytokines, ROS as well as mROS levels during infection. We found that during *S*.*uberis* infection, it is TLR2 deficiency that induced more severe pathological changes through MyD88-mediated signaling pathways. In addition, our work demonstrates that mROS mediated by TLR2 has an important role in host defense response to combat *S. uberis* infection in mammary glands as well as MECs.

## Introduction

Mastitis is a type of inflammation mainly caused by intramammary infection and causes great harm to the dairy industry [1]. *Streptococcus uberis* is an environmental pathogen that is emerging as the most important mastitis-causing agent in some regions [2]. Previous studies in our laboratory have demonstrated that persistent inflammation including swelling, secretory epithelial cell degeneration, and polymorphonuclear neutrophilic leukocyte (PMN) infiltration occurs in mammary tissue following injection with *S. uberis* [3]. The inflammatory responce caused by *S. uberis* is lighter than that caused by *E. coli* [3].These pathological response connected with intracellular infection of *S. uberis* as it escaped the elimination of immune cells and formed persistent infection.

Activation of pattern recognition receptors (PRRs) to produce natural inflammatory immune response is important to control the intracellular infection induced by bacteria like *S. uberis* [4]. TLR family plays a critical role in these processes. Upon activation by microbes, the MyD88-dependent pathway triggers production of inflammatory cytokines through activation of nuclear factor (NF)-κ B and mitogen-activated protein kinases, and/or a TIR-domain-containing adapter-inducing interferon (IFN)-β (TRIF)-dependent pathway associated with induction of IFNs and stimulation of T cell responses [5]. Previous research has found that PRRs are not only expressed by immune cells, but also in conventional non-immune cells, for example, endothelial and epithelial cells, which also contribute to immune regulation [6].

Strandberg et al. demonstrated for the first time in 2005 that TLRs and their downstream molecules are expressed on bovine MECs [6]. Ibeagha-Awemu et al. further demonstrated that expression of TLR4, MyD88, NF-κB, TIR domain-containing adapter molecule 2 (TICAM2) and IFN-regulatory factor 3 increased in bovine MECs challenged by lipopolysaccharide [7]. These studies announced that MECs might have a pivotal role in host defense with TLRs for their huge number in the mammary gland. Our laboratory did a lot of work on the function of TLRs and MECs in eliminating *S. uberis* infection *in vivo* and *in vitro* models. We found that TLRs, mainly TLR2 but no not excluding TLR4, initiated a complex signaling network characterized by NF-κ B and nuclear factor of activated T cells. In addition, they activated the secretion of cytokines and chemokines accompanied with their self-regulation pathways in response to *S. uberis* challenge.

Reactive oxygen species (ROS) are free radicals that contain oxygen atoms, including hydrogen peroxide (H_2_O_2_), superoxide anion (O^2 −^) and hydroxyl radical (OH^−^) [8]. They are produced intracellularly through multiple mechanisms depending on the cell and tissue types. However, the two major sources in mammalian cells are membrane-associated NADPH oxidase-induced and the mitochondria [9]. It has been reported that, in most tissues, mROS from the respiratory chain are important [10]. Mitochondria function as a defense against bacterial infection in innate immunity, mainly through mROS, which is demonstrated by the fact that mROS modulate several signaling pathways, including NF-κ B, C-Jun N-terminal kinase and the caspase-1 inflammasome [11]. Previous studies have shown that restriction of pathogen-induced mROS impairs NF-κB activation, suggesting that mROS positively control the NF-κB signaling pathway [12, 13]. In addition, the production of mROS in immune cells (e.g. macrophages) involves recruitment of tumor necrosis factor (TNF) receptor-associated factor (TRAF)6 to mitochondria, which also acts as an adaptor of the TLR signaling pathway [14]. MECs are the main cells for lactation in mammary tissue. In recent years, they have also been found to play a non-negligible role in the regulation of infection. Pathogens invading mammary tissue and epithelial cells can stimulate MECs to produce proinflammatory cytokines, anti-inflammatory factors and chemokines such as TNF-α, interleukin (IL)-1β, IL-4, IL-6, IL-8 and IL-10 [15]. It is possible that MECs are also involved in the generation of ROS in infection. However, few studies have investigated the interaction between TLRs and mROS against *S. uberis* infection *in vitro and in vivo*. Therefore, we investigated whether the process of TLR induction of mROS production plays an important role against *S. uberis* infection in host and MECs.

## Results

### TLR2 mediates tissue damage and anti-*S. uberis* infection in mammary glands

*S. uberis* belongs to gram-positive bacteria which is mainly recognized by TLR2. However, previous research had demonstrated that the role of TLR4 could not be ignored in *S. uberis* infection for their close relationship, similar structure and function [16, 17]. In this work, we explored the role of TLR2 and TLR4 in *S. uberis* infection in TLR2^−/−^ and TLR4^−/−^ mice to understand further the molecular defence mechanisms in *S. uberis* mastitis. No histological changes were observed in WT-B6 or WT-B10 mammary glands of control mice, whereas, there was some suspicion of tissue damage in TLR2^−/−^ and TLR4^−/−^ control mice (Fig 1A). Inflammation and tissue damage appeared in mammary tissue after infection with *S. uberis* in all challenged groups. This response was characterized by PMN infiltration, increased bleeding and epithelial cell degeneration, and excess adipose tissue. Compared with WT-B6 mice, TLR2 deficiency induced more severe pathological damage. A higher score was present for the three indexes mentioned above and there were significant increases in bleeding and degeneration, and excess adipose tissue (*P* < 0.05; Fig 1B). However, TLR4 deficiency caused no inflammation and tissue damage during *S. uberis* challenge (Fig 1A and 1C).

**Fig 1.**
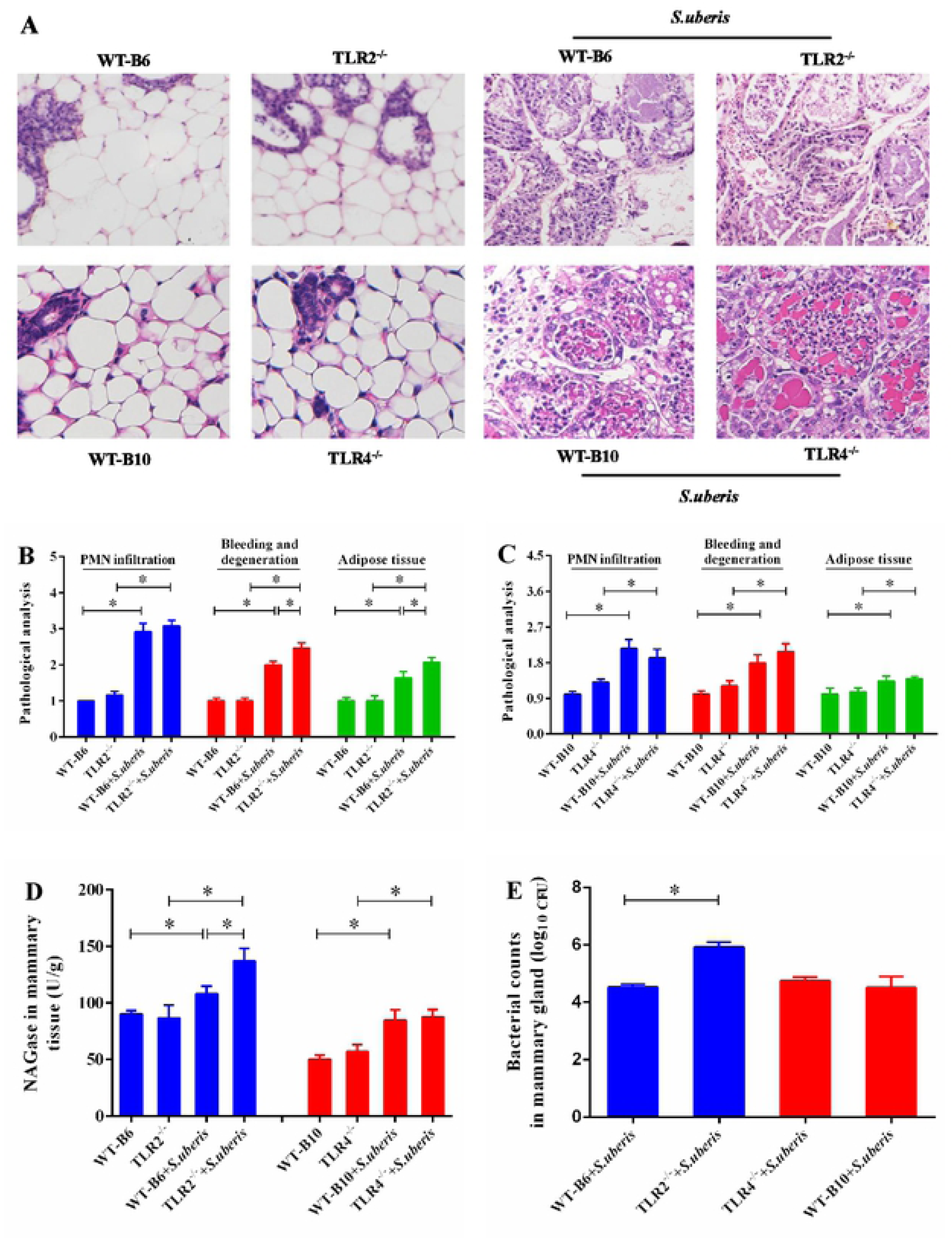
TLR2 mediates tissue damage and anti-S. *uberis* infection in mammary gland. (A, B, C) Mammary glands were stained through hematoxylin and eosin, and the bleeding and degeneration, PMN infiltration, adipose tissue were analyzed by light microscopic. (D) NAGase activity was analyzed in mammary glands. (E) Viable bacteria was counted via the plate with THB agar medium. Data are presented as the means ± SEM (n=6). \**(P*< 0.05) = significantly different between the indicated groups.

NAGase, a marker enzyme of MECs and mammary gland damage, was significantly elevated in TLR2^−/−^ mice, but not in TLR4^−/−^ mice when compared with WT mice at 24 h post-challenge (*P* < 0.05; Fig 1D). Similarly, the bacterial numbers were higher in the mammary tissue of TLR2^−/−^ mice than in WT-B6 mice (*P* < 0.05) but there was no significant difference between TLR4 ^− / −^ and WT-B10 mice. We conclude that TLR2 primarily mediated the tissue damage and antibacterial effect in mammary glands during *S. uberis* infection.

### TLR2 and TLR4 deficiency affect secretion of cytokines in *S*.***uberis* infection**

The secretion of proinflammatory cytokines in mammary glands which better reflects the level of inflammation have been reported previously [18, 19]. However, the massive release of cytokines can cause irreversible damage to tissues. Here, we investigated the role of TNF-α, IL-1β and IL-6 in response to *S. uberis* infection in TLR2^−/−^ and TLR4^−/−^ mice (Fig 2A and 2B). *S. uberis* challenge increased TNF-α level significantly in WT, TLR2 ^− / −^ and TLR4 ^− / −^ mice (*P* < 0.05). Compared with corresponding WT mice, TNF-α and IL-1β in TLR2^−/−^ mice and TNF-α in TLR4^−/−^ mice significantly decreased (*P* < 0.05). These results indicate that TLR2 and TLR4 deficiency affected secretion of cytokines in *S*.*uberis* infection.

**Fig 2.**
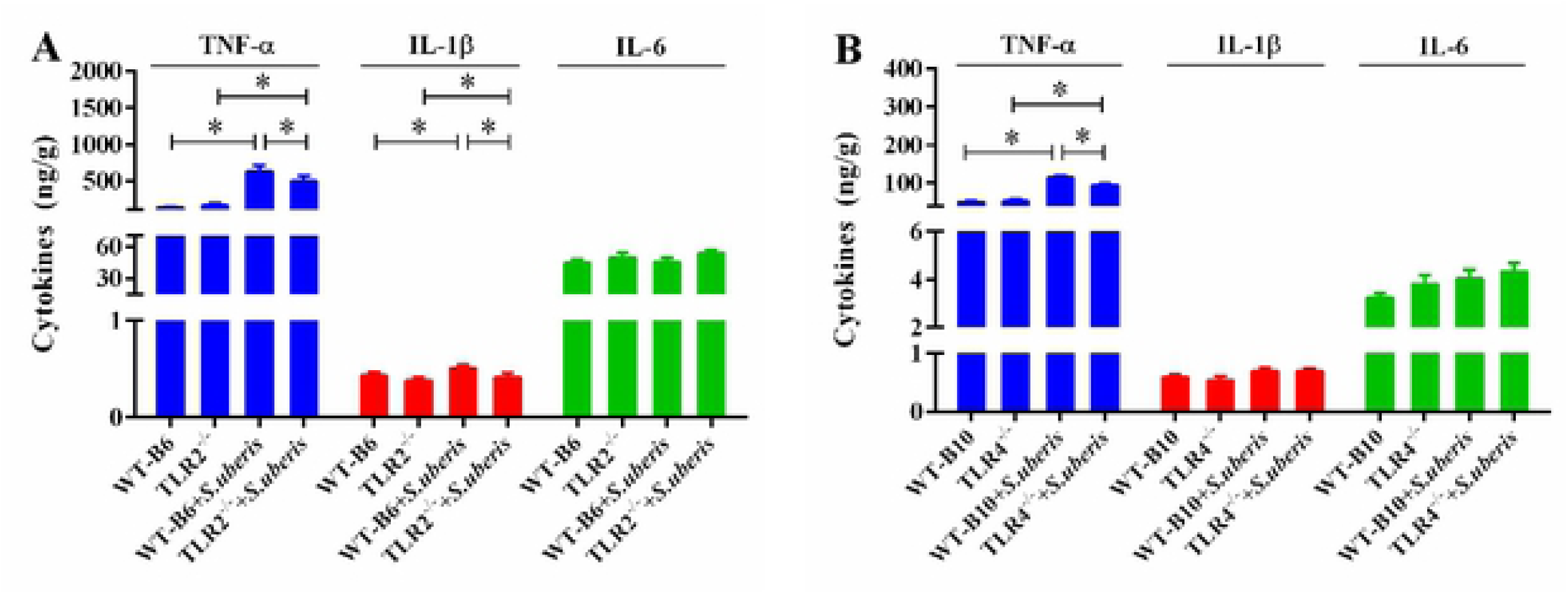
TLR2 and TLR4 deficiency affect the secretion of cytokines in *S*.*uberis* infection. (A, B) The protein expression of TNF-α, IL-I ß and IL-6 were determined by ELISA in mammary glands. Data arc presented as the means ± SEM (n= 6). * (*P*< 0.05) = significantly different between the indicated groups.

### MyD88-dependent pathway predominates in *S. uberis* infection

MyD88 dependent and independent pathways are vital in host’s response after TLRs activated [20, 21]. We assessed the expression of MyD88 and TRIF respectively by immunohistochemistry in mammary glands, which are the key molecules of the MyD88-dependent or independent signaling pathways. There was a significant increase in MyD88 but not TRIF expression in WT, TLR2^−/−^ and TLR4^−/−^ mice (*P* < 0.05). Lower levels of MyD88 were observed in TLR2^−/−^ and TLR4^−/−^ mice compared with WT mice (*P* < 0.05; Fig 3A and 3B). As the main functional cells in mammary glands, our former study had established that MECs played a key role in anti-infection response in mammary glands. We also detected that interference of TLR2 and/or TLR4 by specific siRNA significantly decreased MyD88 expression in *S. uberis* infection (*P* < 0.05; Fig 3C, 3D and 3E). These data suggest that the MyD88-dependent pathway predominates in *S. uberis* infection in MECs and in mammary glands following TLR activation.

**Fig 3.**
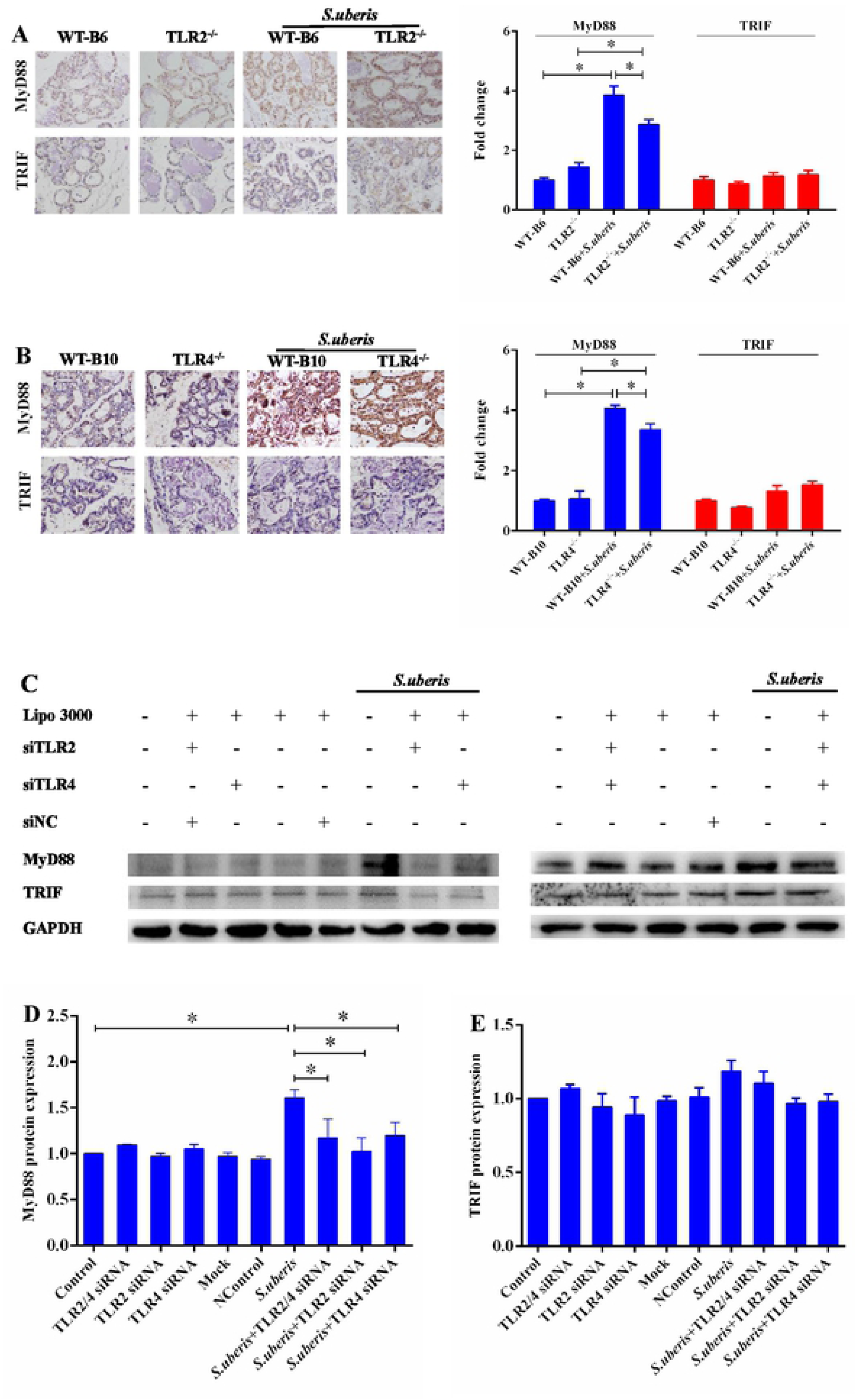
MyD88 dependent pathway predominates in *S. uberis* infection. (A, B) Immunohistochemistry was used to analyze the expression of MyD88 and TRIF in mammary glands. Data are presented as the means ± SEM (n= 6). * (*P*< 0.05) = significantly different between the indicated groups. (C, D, E) The protein expression of MyD88 and TRIF were determined by Western blot in MECs. Data arc presented as the means ± SEM (n= 3). * (*P*< 0.05) = significantly different between the indicated groups.

### TRAF6 and ECSIT participate in signal sensing from TLRs in *S. uberis* infection

We next evaluated the level of TRAF6 and ECSIT, which are downstream targets of the MyD88 signaling pathway, using immunohistochemistry in mammary glands of mice. The adaptors TRAF6 and ECSIT increased dramatically in all mice after *S. uberis* infection (*P* < 0.05), although TLR2 or TLR4 deletion weakened expression of ECSIT in WT compared with TLR2 ^− / −^ and TLR4 ^− / −^ mice (*P* < 0.05; Fig 4A and 4B). In MECs, interference of TLR2 or TLR4 significantly reduced expression of TRAF6 and ECSIT after *S. uberis* infection (*P* < 0.05; Fig 4C, 4D and 4E). TRAF6 and ECSIT expression did not differ between TLR2 ^− / −^ and TLR4 ^− / −^ mice. The results confirm that TRAF6 and ECSIT downstream of the MYD88 pathway mediate the anti-*S. uberis* response in mice and MECs.

**Fig 4.**
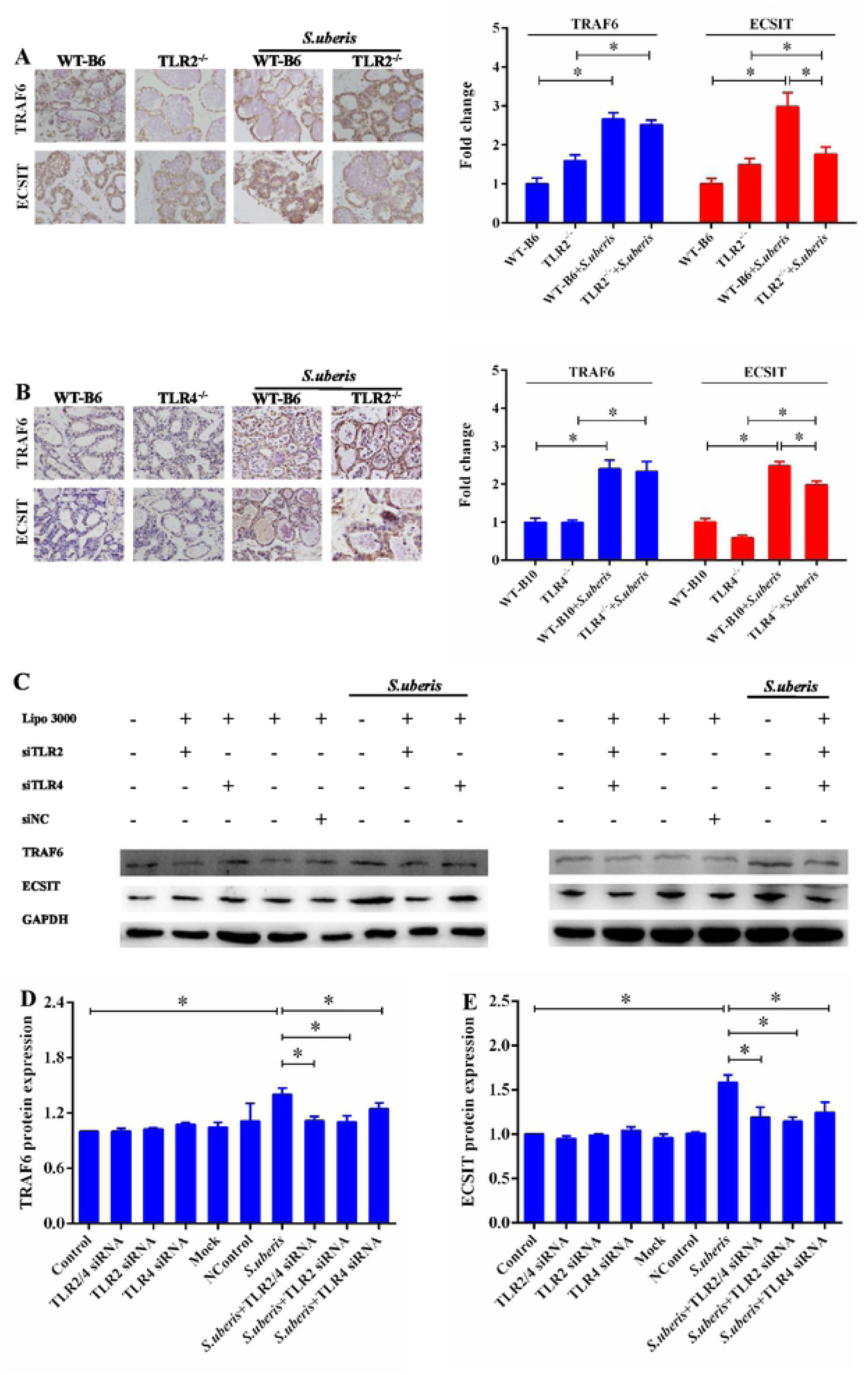
TRAF6 and ECSIT participate in scnseing signal from TLRs in *S. uberis* infection. (A, B) lmmunohistochemistry was used to analyze the expression ofTRAF6 and ECSIT in mammary glands. Data are presented as the means ± SEM (n= 6). \**(P*< 0.05) = significantly different between the indicated groups. (C, D, E) The protein expression of TRAF6 and ECSIT were determined by Western blot in MECs. Data arc presented as the means ± SEM (n= 3). \**(P*< 0.05) = significantly different between the indicated groups.

### TLRs mediate redox status of mammary glands during *S. uberis* infection

TRAF6 activated by TLRs transfers from the cytoplasm to mitochondria, where it engages ECSIT to produce mROS, which induces cellular anti-bacterial response [22]. Since the level of ROS in tissue cannot be detected well, we analyzed T-AOC, SOD, MDA and UCP2 in mammary glands to reflect indirectly the antioxidant levels. The levels of MDA and UCP2 were significantly increased due to the infection of *S. uberis* in WT and TLR2 ^− / −^ and TLR4 ^− / −^ (*P* < 0.05; Fig 5A and 5B). There was no obvious distinction between TLR2^−/−^, TLR4^−/−^ and WT mice (*P* > 0.05). T-AOC was significantly lower in all groups of mice after *S. uberis* challenge (*P* < 0.05). Deletion of TLR4 rather than TLR2 significantly decreased SOD activity after *S. uberis* infection (*P* < 0.05). These results indicate that the host’s oxidation level does change after *S. uberis* infection and these changes are related to the TLR signaling pathway.

**Fig 5.**
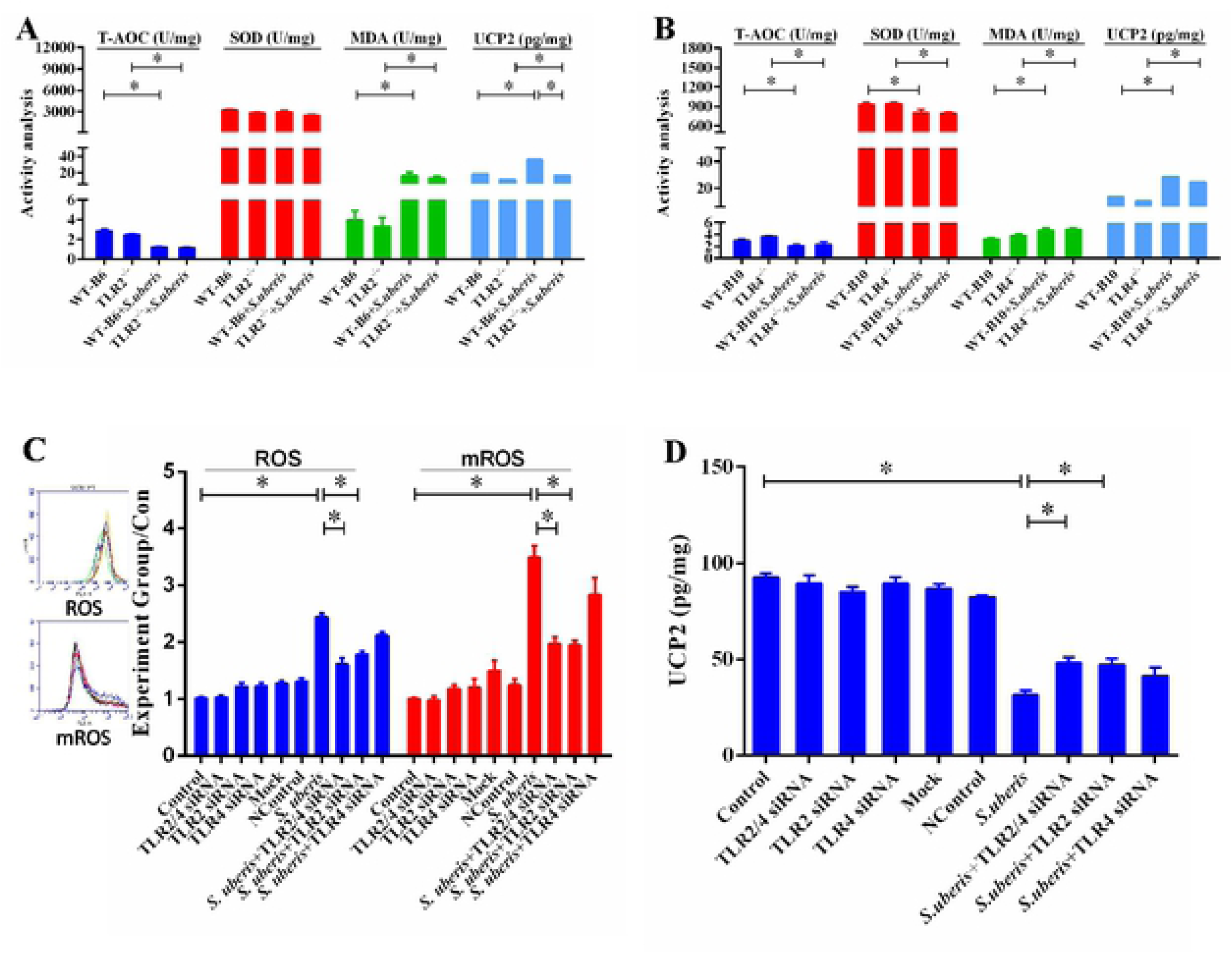
TLRs mediate redox status of mammary glands during *S. uberis* infection. (A, B) The protein expression of T-AOC, SOD, MDA and UCP2 were determined by kits in mammary glands. Data are presented as the means ± SEM (n=6). \**(P*< 0.05) = signiificantly different between the indicated groups. (C) CcllQuest Pro acquisition and analysis software analy1.cd ROS and mROS in MECs. (D) The activity of UCP2 were determined by ELISA in MECs. Data are presented as the means± SEM (n= 3). *(*P*< 0.05) = significantly different between the indicated groups.

We aimed to clarify whether MECs were involved in the change in redox status and had a crucial role in *S. uberis* infection after activation of the TLR signaling pathway. We interfered with the expression of TLR2 and/or TLR4 in EpH_4_-Ev cells and detected ROS, mROS and UCP2 levels. *S. uberis* infection caused a significant increase in ROS and mROS (Fig 5C and 5D). Special targeting of siRNA to TLR2 significantly reduced the level of ROS and mROS (*P* < 0.05). SiTLR4 decreased their levels to some extent, but no significant difference was observed (*P* > 0.05). Expression of UCP2 decreased, and interference with siTLR2 reversed this change (*P* < 0.05). Taken together, these results demonstrate that infection with *S. uberis* changed the redox status of mammary glands and MECs, and TLR2 played an essential role in this process, especially in MECs.

### mROS play an important role against *S. uberis* infection in MECs

GKT137831, a specific inhibitor of NADPH oxidase 1 (NOX1) and NOX4; and NG25, an inhibitor of TAK1 [23, 24], were used to suppress ROS generation from NOX complexes and to down-regulate production of proinflammatory cytokines, respectively. GKT137831 and NG25, simultaneously or separately, reduced the generation of ROS but not mROS after challenge with *S. uberis* (*P* < 0.05; Fig 6A). The bacterial counts of *S. uberis* in MECs were significantly higher in the inhibitor-treated groups (*P* < 0.05) (Fig 6B). We inhibited production of mROS by siECSIT to establish mROS role in regulating inflammation and anti-*S. uberis* activity. ROS and mROS levels decreased significantly after using siECSIT (*P* < 0.05; Fig 6D). Similar results were observed for TNF-α, IL-1β and IL-6 expression; their levels were up-regulated after *S. uberis* infection while siECSIT reduced them (Fig 6E). The bacterial counts of *S. uberis* in MECs were significantly higher in the siECSIT treatment group (Fig 6F). These results demonstrate that mROS does play an important role against *S. uberis* infection in MECs.

**Fig 6.**
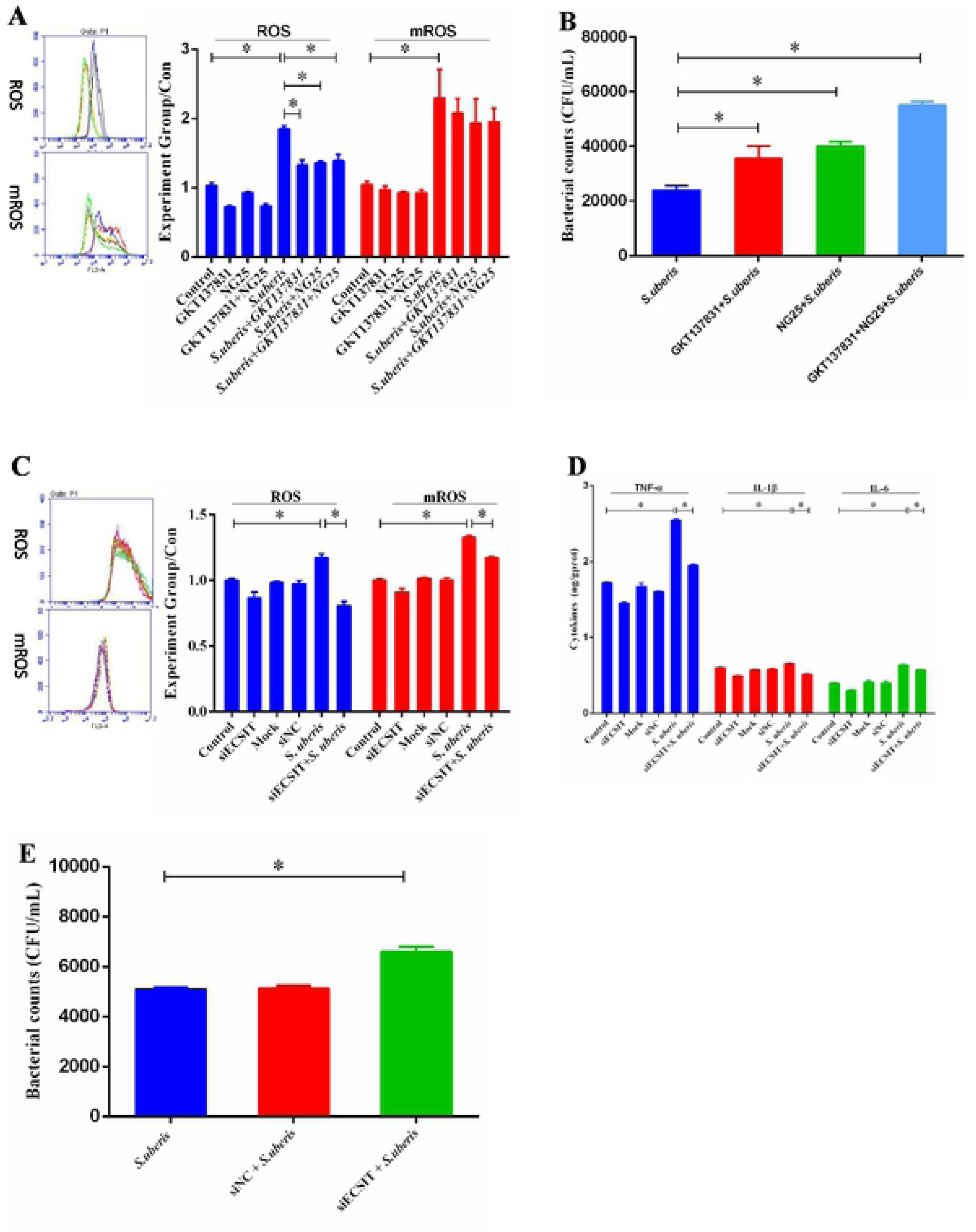
mROS play an important role in anti-*S. uberis* infection in MECs. (A) The expression of ROS and mROS after using GKTl3783 I and NG25 simultaneously or separately during *S. uberis* infection in MECs. (B) Viable bacteria was counted via the plate with THB agar medium after using GKT13783 l and NG25 simultaneously or separately during *S. uberis* infection in MECs. (C) The expression of ROS and mROS after using siECS!T in MECs. (D) The expression of TNF-α, IL-Iβ and IL-6 after using siECSIT in MECs. (E) Bacteria counts after using siECSIT in MECs. Data arc presented as the means ± SEM (n= 3). \**(P* < 0.05) = significantly different between the indicated groups.

## Discussion

The intrusion signal (from molecules broadly shared by pathogens that could be recognized by the immune system) of intracellular bacteria captured by PRRs is crucial for host control of inflammation and pathogen proliferation [25]. TLRs are one of the most ancient, conserved components of the immune system, and have been established by our laboratory to sense and respond to *S. uberis* [3]. *S. uberis* is a kind of Gram-positive bacterium and TLR2 is the principal receptor that can sense its invasion [26]. However, TLR2 and TLR4 share the same delivery system, and current studies have not yet distinguished their exact roles in defending against *S. uberis* infection. We used TLR2^−/−^ and TLR4^−/−^ mice to investigate, for the first time thoroughly, the roles of these two high-correlation receptors in *S. uberis* infection. Deficiency of TLR2, but not TLR4, induced a more severe inflammatory response and tissues damage in mammary gland and bacterial viability was higher. These results confirmed that TLR2 detected *S. uberis* infection, initiated the antibacterial immunological reaction and controlled the inflammatory status in mammary glands.

Proinflammatory cytokines, such as TNF-α, IL-1β and IL-6, are secreted following activation of TLRs and their respective downstream signaling pathways mainly in immune cells [27]. They are involved in upregulation of inflammatory reactions and play a role in regulating host defense against pathogens mediating the innate immune response. In this study, TNF-α, the initiating factor in the cytokine storm, increased dramatically after *S. uberis* challenging in variant mice. This change was only seen in TLR2^−/−^ mice for IL-1β. No changes were observed for IL-6. These findings were consistent with previous reports that expression of TNF-α, IL-1β and IL-6 had a chronological order, and for the samples detected here, were only expressed at 24 h post-infection [28]. Compared with WT mice, in TLR2^−/−^ mice, TNF-α and IL-1β were obviously decreased. This further demonstrated the important role of TLR2 in the interaction of *S. uberis* infection with the host. TNF-α levels in TLR4 ^− / −^ mice had similar changes. This could be explained by the fact that inflammatory response networks are complex after infection. Positive and negative inflammatory factor feedback loops both exist in *S. uberis*-infected mammary glands. The secondary inflammation induced by initial inflammatory factors might result partly from activation of TLR4. Hence, down-regulation of TNF-α caused by deficiency of TLR4 was not the same as that losing TLR2, which could not neutralize inflammatory response challenged by *S. uberis*.

Two distinct signaling pathways, the MyD88-dependent and TRIF-dependent pathways, are triggered by dimerized and activated TLRs [29]. Our experiments *in vivo* found that MyD88 instead of TRIF was affected significantly in *S. uberis* infection, and thus confirmed that the MyD88-dependent pathway predominated in this process. This phenomenon also existed in other bacterial infections. For example, Wiersinga et al. reported it in *Burkholderia pseudomallei* infection [30]. In the MyD88-dependent pathway, MyD88 recruits IL-1 receptor-associated kinases and then phosphorylates and activates TRAF6, which in turn polyubiquinates TAK1, and induces secretion of inflammatory cytokines in our research on *S. uberis* infection *in vivo* and *in vitro* [3, 31]. Recently, it have been suggested that activated TRAF6 translocates to the mitochondria, which leads to ECSIT ubiquitination, resulting in increased mROS generation [32]. This signaling pathway plays an important role in the innate immune response against intracellular bacteria. A recent study has also shown that ECSIT-and TRAF6-depleted macrophages have decreased levels of TLR-induced ROS and are significantly impaired in their ability to kill intracellular bacteria [13]. Sonoda et al. similarly found that estrogen-related receptor α and PPAR gamma Coactivator-1 β (PGC-1ß) act together as key effectors of IFN-γ-induced mitochondrial ROS production and host defense [33]. Our study emphasizes the importance of mROS in killing bacteria. Since accumulating ROS in tissues is difficult, previous research has always detected the presence of members of the antioxidant system, such as T-AOC, SOD, MDA and UCP2, in organs to reflect production of ROS indirectly [34, 35]. Our results showed that *S. uberis* challenge caused changes in redox status of mammary glands. This indicates that the TLRs/MyD88/TRAF6/ECSIT/mROS axis participated in the defense response to *S. uberis* infection.

The inflammatory phenomena from mammary glands involve integrated responses of all kinds of mammary cells including macrophages, PMNs, lymphocytes, MECs and even matrix cells [36]. In the past decade, we paid more attention to the defensive ability of MECs because they are the most numerous cells in the udder, and we have detected TLRs-mediated signaling pathways and secretion of more than 40 cytokines [28]. In addition, we showed that *S. uberis* adhered and internalized in MECs, which establishes that MECs are one of the main target cells of *S. uberis* (data not published). Intriguingly, MECs are not real immune cells and have a distinctive response to bacterial infection. For example, we have found that the PI3K/Akt/mTOR pathway in MECs generates a positive contribution to inflammation following viable *S. uberis* challenge, which is not consistent with the usual situation in some immune cells [28]. Thus, we treated MECs with specific siRNAs targeting to TLR2 and/or TLR4 and then evaluated the effect of *S. uberis* challenge on the expression of key adaptor proteins. The data confirmed that the MyD88-dependent pathway predominated in *S. uberis*-infected MECs after TLR2 activation. A similar signal transfer process was reported in macrophages infected with *Mycobacterium tuberculosis*, another Gram-positive bacterium [37]. In the present study, we were interested in whether the TLR2/MyD88/TRAF6/ECSIT axis regulated production of ROS. Suppression of TLR2, but not TLR4, reduced the level of ROS and mROS in MECs after *S. uberis* challenge. This result was further explained by the detection of UCP2, which separates oxidative phosphorylation from ATP synthesis and thus improves the production of ROS and mROS [38].

Initially, mROS was considered to be a by-product of bio-oxidation, and its synthesis cannot be regulated. A large body of researches have established that oxidative phosphorylation in mitochondria is the main pathway for mROS production and is the main source of ROS [39]. To express catalase in mitochondria could effectively reduce the production of mROS, thereby reducing the killing effect of macrophages on pathogens, indicating that mROS is a key driver in the process of antibacterial activity [40]. Our current study showed that TLR2 regulates the generation of ROS, including mROS, during *S. uberis* infection both *in vivo* and *in vitro* [3]. We suggest that TLR2-mediated mROS are involved in *S. uberis* infection. To confirm our hypothesis, GKT137831 and NG25 were used alone or simultaneously to suppress ROS from NOX complexes and production of proinflammatory cytokines. We found that the antibacterial activity of MECs was restrained to some extent, and this established that ROS from NOX complexes and cytokines were involved in the host defensive reaction. This is consistent with our previous study [28]. Furthermore, we inhibited the synthesis of mROS by siECSIT and investigated the changes in inflammation and the effect of reducing mROS on bacterial viability. The bacterial counts of *S. uberis* were significantly higher in the siECSIT treatment group. These results demonstrate that TLR2-mediated mROS are a key factor against *S. uberis* infection in MECs.

In conclusion, mROS participate in the host response against *S. uberis* infection, and TLR2 is involved in sensing *S. uberis* invasion and controlling mROS production by regulating expression of TRAF6 and ECSIT. Additionally, the function of mROS against *S. uberis* infection probably relies on their ability to regulate cytokine levels, thereby controlling the level of inflammation. This study increased our understanding of the molecular defense mechanisms in *S. uberis* mastitis, and provides theoretical support for the development of prophylactic strategies for this critical disease.

## Materials and methods

### Bacterial strain, cell culture and treatment

*S. uberis* 0140J (American Type Culture Collection, Manassas, VA, USA) was inoculated into Todd–Hewitt broth (THB) supplemented with 2% fetal bovine serum (FBS; Gibco, USA) at 37°C in an orbital shaker to mid-log phase (OD_600_ 0.4–0.6). MECs (American Type Culture Collection) were incubated in Dulbecco’s modified Eagle’s medium (DMEM) with 10% FBS and plated at 80% confluence in 6-well cell cultrue cluster. After culture in serum-free DMEM for 4 h, the monolayer was treated with 40 nM NG25 (inhibitor of TGFβ-activated kinase 1; TAK1: Invitrogen, Carlsbad, CA, USA) for 24 h; 4 µm MK2206 (inhibitor of NADPHase: SellecK Chemicals, Houston, TX, USA) for 24 h; or transfected with 50 nM siTLR2 or/and siTLR4 for 72 h. SiECSIT with 20 nM were performed for 48 h using Lipofectamine 3000 reagent (Invitrogen). The sequences of siRNA were designed and listed as follows. siTLR2: GTCCAGCAGAATCAATACA; siTLR4: CAATCTGACGAACCTAGTA; siECSIT: GGTTCACCCGATTCAAGAA. The treated cells were infected with *S. uberis* at a multiplicity of infection (MOI) of 10 for 2 or 3 h at 37°C. The supernatant and cells were collected separately and stored at -80 °C until use.

### Mice and treatment

Mice, including wild-type C57BL/6 (WT-B6), wild-type C57BL/10 (WT-B10), TLR2^−/−^ (C57BL/6) and TLR4^−/−^ (C57BL/10), aged 6–8 weeks were purchased from Nanjing Biomedical Research Institute of Nanjing University (Nanjing, China) and bred under specific pathogen-free conditions in the Nanjing Agricultural University Laboratory Animal Center. All experimental protocols were approved by the Regional Animal Ethics Committee and in compliance with Animal Welfare Act regulations as well as the Guide for the Care and Use of Laboratory Animals.

Seventy-two hours after parturition, all experimental groups of female mice were infused with 50 μL *S. uberis* according to the number of complex infections (MOI = 10) into the left 4 (L4) and right 4 (R4) teats. The animals in the control groups were infused with same volume of phosphate-buffered saline (PBS). The offspring were weaned 1 h prior to experimental infusion. All mice were killed 24 h post-infusion. The mammary glands and serum were aseptically collected and stored at −80°C.

Mammary gland was fixed in 10% neutral buffered formalin. Sections of 5 μm thickness were stained with hematoxylin and eosin. Mammary gland tissues were weighed and homogenized with sterile PBS (1:5, W/V) on ice. After centrifuged at 500 *g* at 4°C for 40 min, the supernatant was centrifuged again. The second supernatant was collected and stored at −80°C until assayed.

### Histological observation and immunohistochemistry

The mammary tissue fixed in 10% neutral buffered formalin was trimmed and flushed in water for at least for 4 h, and then dehydrated in alcohol solutions ranging from 75% to 100%, with 5% increase at 1 h intervals. After soaking in xylene, the tissues were embedded in wax for 3 h at 60°C. Slices (5 μm thick) were cut and stained with hematoxylin and eosin. The histological changes including PMN infiltration, bleeding and degeneration, and adipose tissue loss were analyzed by light microscopy (BH2; Olympus, Tokyo, Japan) at a magnification of 40×. Four sections of mammary tissue were quantified for each animal. Ten fields were randomly selected per tissue section and assigned a score of 1, 2 or 3 based on the degree of damage.

Immunohistochemical staining was performed as follows. Tissue sections were washed with PBS, then covered with 3% H_2_O_2_ for 15 minutes at 37°C to inhibit further endogenous peroxidase activity. Tissue slices were blocked with 5% bovine serum albumin and incubated with antibodies against MyD88, TRAF6, ECSIT and TRIF (Cell Signaling Technology, Danvers, MA, USA), at 4°C in a humidified chamber. Overnight, biotinylated anti-rabbit IgG (Boster Bio-Technology, Wuhan, China) was incubated for 30 min at 37°C. After rehydration, the sections were incubated with avidin–biotin peroxidase complex for 40 min at 37°C. Finally, the sections were washed and bound conjugates were revealed by diaminobenzidine staining (Boster Bio-Technology).

### RNA extraction and quantitative real-time polymerase chain reaction (PCR)

PCR was carried out as previously described [41]. Total RNA was extracted by TRIzol reagent (TaKaRa, Dalian, China). Corresponding cDNA was obtained using reverse transcriptase and Oligo (dT) 18 primer (TaKaRa). An aliquot of the cDNA was mixed with 25 µL SYBR® Green PCR Master Mix (TaKaRa) and 10 pmol of each specific forward and reverse primer. All mixed systems were analyzed in an ABI Prism 7300 Sequence Detection System (Applied Biosystems, Waltham, MA, USA). Fold changes were calculated as 2 ^− ΔΔCt^. All primer sequences (Table S1) were synthesized by Invitrogen Biological Company (Shanghai, China).

### Total protein extraction and western blotting

Cells were washed twice in 2 mL ice-cold PBS and collected in an Eppendorf tube (gently scraped by a rubber policeman) after being lysed on ice for 20 min in lysis buffer (Beyotime, Nantong, China). Extracts with equal amounts of proteins were solubilized by SDS sample buffer (BioRad, Califonia, USA), separated by SDS-PAGE, and transferred to polyvinylidene difluoride membranes (Millipore, Bedford, MA, USA). The membranes were incubated with corresponding polyclonal antibodies: anti-MyD88, anti-TRAF6, anti-ECSIT anti-TRIF, and anti-GAPDH. The signals were detected by an ECL western blot analysis system (Tanon, Shanghai, China). Analysis of bands was quantified with Image J software (NIH, Bethesda, MD, USA).

### Measurement of ROS and mROS

EpH4-Ev cells were incubated in dichloro-dihydrofluorescein diacetate (10 μM, 30 min) (Beyotime, Nantong, China) or MitoSOX (5 μM, 20 min) (Thermo, Waltham, MA, USA) at 37°C, washed three times in PBS and detached. The cells were centrifuged at 400 *g* for 5 min, resuspended in PBS, and immediately analyzed by flow cytometry using FACSCanto (BD, New Jersey, USA). Ten thousand cells per sample were analyzed using CellQuest Pro acquisition and analysis software.

### Assay of TNF-α, IL-1, **and IL-6 by ELISA**

The levels of TNF-α, IL-1 β, and IL-6 in mammary glands and EpH_4_-Ev cells were measured by ELISA (Rigor Bioscience, Beijing, China). Prepared standards (50 μL), and antibodies (40 μL) labeled with enzyme (10 μL) were reacted for 60 min at 37°C and the plate was washed five times. Chromogen solutions A (50 μL) and B (50 μL) were added and incubated for 10 min at 37°C. Stop solution (50 μL) was added and optical density value was measured at 450 nm within 10 min. Qualitative differences or similarities between the control and experimental groups were consistent throughout the study.

### Detection of NAGase, T-AOC, SOD, MDA and UCP2

The activities or levels of *N*-acetyl-β-D-glucosaminidase (NAGase), total antioxidant capacity (T-AOC), superoxide dismutase (SOD), malondialdehyde (MDA) and uncoupling protein 2 (UCP2) were determined using commercial kits purchased from Nanjing Jiancheng Bioengineering Institute (China).

### Viable bacterial count assay

Viable bacteria were enumerated as colony-forming units (CFU) on THB agar. The mammary glands were aseptically homogenized with sterile PBS (1:5, W/V). The supernatants were spread on plates. CFUs were counted by the spread plate method after incubation for 12 h at 37°C.

MECs and MECs with siECSIT were incubated in DMEM with 10% FBS and plated at 80% confluence in 6-well plates. After culture in serum-free DMEM for 4 h, at mid-exponential phase (OD_600_ 0.4–0.6), *S. uberis-*infected cells were washed 3 times with PBS containing 100 mg/mL gentamicin, followed by gentamicin-free PBS. Cells were pelleted at 1.4 *g* for 10 min. The same number of cells were lysed with sterile triple distilled water, and CFUs were counted by the spread plate method after incubation for 12 h at 37°C.

### Statistical analysis

Results were analyzed using GraphPad Prism 5.0 software (GraphPad Software Inc., La Jolla, CA, USA). Data were expressed as means standard error of the mean (SEM). Differences were evaluated by one-way analysis of variance followed by post-hoc tests. Significant differences were considered at *P* < 0.05.

## Supporting information

**S1 Table. Oligonucleotide sequences used for RT-qPCR**.

**S1 Fig. TLR2/4 mediates the NAGase activity after challenge with *S. uberis* in the supernatant of MECs**.

**S2 Fig. TLR2/4 mediates the inflammatory response after challenge with *S. uberis* in MECs**.

**S3 Fig. The suppression of mROS reduces the inflammation factors after challenge with *S. uberis* in the supernatant of MECs**.

**S4 Fig. The protein expression of ECSIT were determined by Western blot after using siECSIT in MECs**.

## Acknowledgments

This project was supported by grants from the National Natural Science Foundation of China (No. 31672515), the Key Project of Inter - governmental International Scientific and Technological Innovation Cooperation (No.2018YFE0102200), and the Project Funded by the Priority Academic Program Development of Jiangsu Higher Education Institutions. We thank International Science Editing (http://www.internationalscienceediting.com) for editing this manuscript.

## Author Contributions

Conceived and designed the experiments: Bin Li, Zhixin Wan. Performed the experiments: Bin Li, Zhixin Wan, Zhenglei Wang. Analyzed the data: Bin Li, Zhixin Wan, Jiakun Zuo, Yuanyuan Xu. Contributed reagents/materials/analysis tools: Xiangan Han, Vanhnaseng Phouthapane, Jinfeng Miao. Wrote the paper: Bin Li, Zhixin Wan, Jinfeng Miao.

